# Lineage labeling with zebrafish *hand2* Cre and CreERT2 recombinase CRISPR knock-ins

**DOI:** 10.1101/2024.12.04.626907

**Authors:** Zhitao Ming, Fang Liu, Hannah R. Moran, Robert L. Lalonde, Megan Adams, Nicole K. Restrepo, Parnal Joshi, Stephen C. Ekker, Karl J. Clark, Iddo Friedberg, Saulius Sumanas, Chunyue Yin, Christian Mosimann, Jeffrey J. Essner, Maura McGrail

## Abstract

**Background:** The ability to generate endogenous Cre recombinase drivers using CRISPR- Cas9 knock-in technology allows lineage tracing, cell type specific gene studies, and *in vivo* validation of inferred developmental trajectories from phenotypic and gene expression analyses. This report describes endogenous zebrafish *hand*2 Cre and CreERT2 drivers generated with GeneWeld CRISPR-Cas9 precision targeted integration.

**Results:** *hand2*-*2A-cre* and *hand2*-*2A-creERT2* knock-ins crossed with ubiquitous *loxP*-based Switch reporters led to broad labeling in expected mesodermal and neural crest-derived lineages in cardiac, pectoral fins, pharyngeal arch, liver, intestine, and mesothelial tissues, as well as enteric neurons. Novel patterns of *hand2* lineage tracing appeared in venous blood vessels. CreERT2 induction at 24 hours reveals late emerging *hand2* progenitors in the 24 – 48 hour embryo contribute to the venous and intestinal vasculature. Induction in 3 dpf larva restricts *hand2* lineage labeling to mesoderm-derived components of the branchial arches, heart, liver and enteric neurons.

**Conclusions:** *hand2* progenitors from the lateral plate mesoderm and ectoderm contribute to numerous lineages in the developing embryo. Later emerging *hand2* progenitors become restricted to a subset of lineages in the larva. The *hand2* Cre and CreERT2 drivers establish critical new tools to investigate *hand2* lineages in zebrafish embryogenesis and larval organogenesis.

**Graphical abstract:** 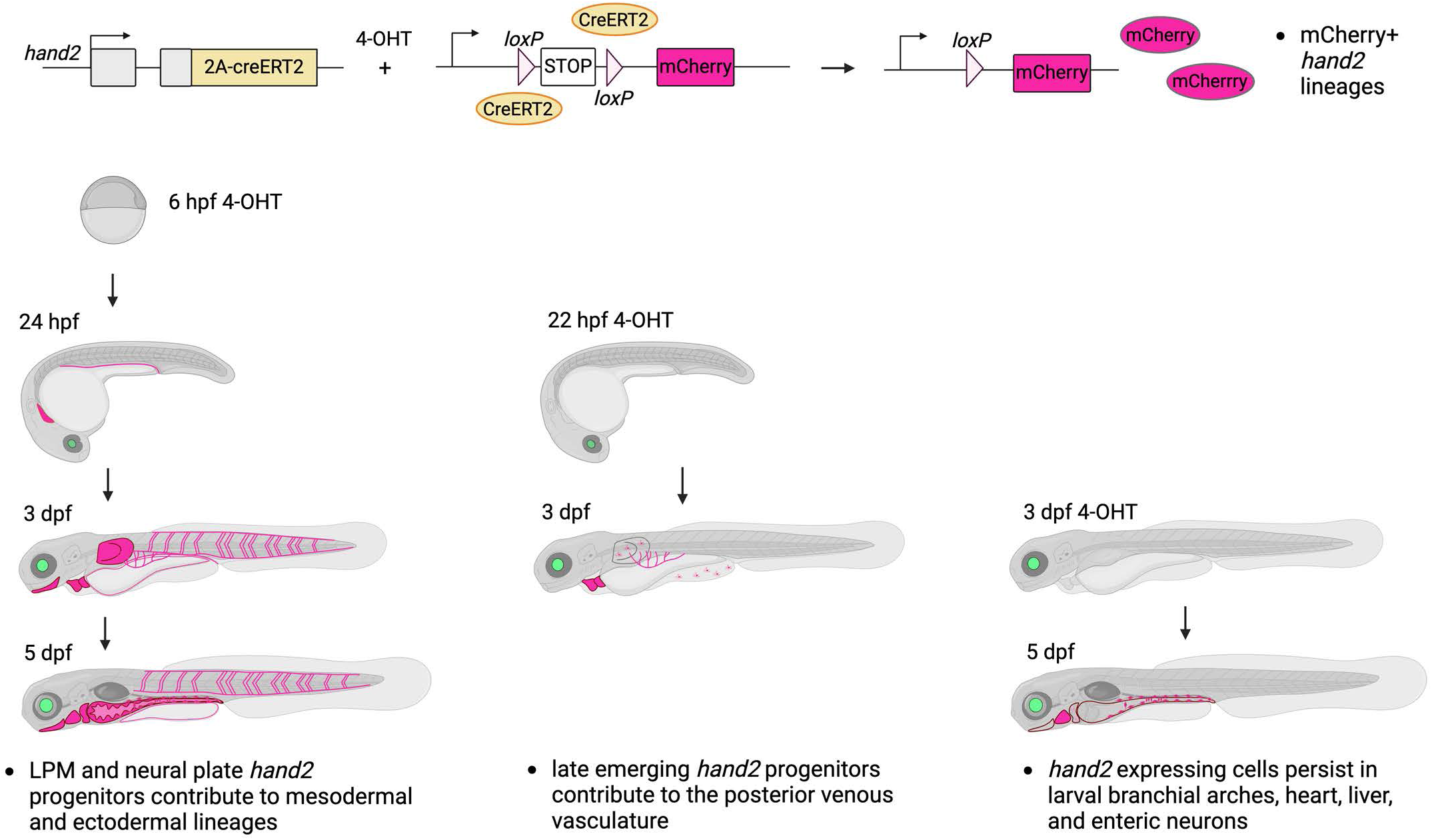

Zebrafish *hand2* Cre and CreERT2 drivers generated with GeneWeld CRISPR/Cas9 precision targeted integration label numerous lineages from the mesoderm and ectoderm. Temporal CreERT2 Tamoxifen regulated switching reveals late emerging *hand2* progenitors contribute to the embryonic venous vasculature and larval organs.

## Introduction

Until recently, methods to create gene-specific lineage reporters and functional genomics tools in zebrafish were limited to molecularly define and clone gene-regulatory elements or enhancer traps screens. With CRISPR-Cas9 knock-in technology, previously inaccessible cell lineages can now be studied by targeted integration of Cre and CreERT2 into gene markers of lineage identity. The GeneWeld CRISPR-Cas9 method for targeted integration directed by short homology ^1,2^ has been successfully applied to generate endogenous Cre drivers in zebrafish proneural genes ^3^ and floxed conditional alleles in cell cycle and epigenetic regulators ^4^. These tools allow robust Cre-mediated proneural-specific conditional gene knockout as well as rescue ^4^, illustrating the power of this approach for cell type specific lineage and gene function studies. Here, as part of our Cre recombinase genetic resource for zebrafish conditional gene studies, we applied GeneWeld to generate Cre and CreERT2 targeted integration lines in the key developmental transcriptional regulator gene *hand2*.

The bHLH transcription factor Hand2 is a key regulator of cell fate specification and differentiation and plays a critical role in organogenesis and the formation of various tissues during development ^5–12^. Hand2 is initially expressed in the embryonic lateral plate mesoderm at the end of gastrulation across vertebrate species ^6–8,13^ and subsequently activated in neural crest cells ^5,9^. Mutant analyses in mice and zebrafish have shown a genetic requirement for Hand2 in development of myocardium, aortic arch, branchial arches, forelimb and pectoral fin mesenchyme, smooth muscle and gut looping, sympathetic neurons, and the posterior vasculature ^9,11,12,14–17^. Recent scRNA-seq profiling of the zebrafish lateral plate mesoderm and lineage tracing in mice identified a conserved role for Hand2 as the earliest marker of mesothelium progenitors ^18,19^. Lineage analysis by persistent reporter expression with a previously described *BAC-based hand2:GFP* reporter ^14^ confirmed Hand2 progenitors contribute to mesothelial tissues in the zebrafish embryo and 3 day post-fertilization larva ^18^. In mice, *Hand2* lineage tracing with defined enhancer-reporter transgenes drive expression in the heart cardiac and neural crest tissues, pharyngeal arches, stomach smooth muscle, limbs and mesothelia ^17,18,20–24^. These tools provide important readouts of *Hand2* expression consistent with its developmental role in transcriptional regulation of mesodermal and neural crest lineages. However, transgenes driving fluorescent reporters that persist prevent analysis of gene expression profiles across timepoints ^20^. To determine the lineage contribution of Hand2-expressing cells during post embryonic development requires inducible lineage drivers that I) allow persistent genetic labeling of cells and ii) fully and faithfully recapitulate endogenous *Hand2* expression, allowing investigation of cell type specific genetic control of development within the *Hand2* lineage.

Here, we describe CRISPR-Cas9 generated *hand2-2A-cre* and *hand2-2A-creERT2* knock-ins in the zebrafish *hand2* gene that provide robust spatial and temporal lineage tracing with ubiquitous floxed reporters. Cre and CreERT2 recombinase activity in the early embryo parallels expected patterns, including *hand2* lineages derived from progenitors in the neural crest that label enteric neurons. *hand2* progenitors in the lateral plate mesoderm contribute to mesodermal components of the branchial arches, heart, pelvic fin and pre-anal fin fold, liver, intestine, and mesothelia. Together our results are consistent with previous analyses of hand2 expression and genetic requirements during organogenesis and tissue development. *hand2-2A-creERT2* Tamoxifen-regulated lineage tracing reveals contributions of early *hand2* progenitors and late emerging-*hand2* expressing cells to the venous vasculature, consistent with the *in vivo* genetic requirement for *hand2* in trunk endothelial development ^12^. Lineage tracing at the 3 dpf larval stage reveals a subset of *hand2* lineages remain active in post-embryonic tissues, reaffirming the continued *hand2* transcriptional activity in larval organogenesis. Our novel tools for *hand2*-based cell type-specific Cre recombination enable studies of lineage contributions and genetic requirements of *hand2* progenitors in embryonic development, post-embryonic organogenesis, and disease modeling.

## Results

### Isolation of *hand2-2A-cre/creERT2* lines by GeneWeld CRISPR targeted integration

To recover endogenous zebrafish *hand2* Cre and CreERT2 drivers, GeneWeld CRISPR-Cas9 targeted integration ^1,2^ was used to knock-in a *2A-cre* and a *2A-creERT2* cassette at the 3’ end of the coding sequence in *hand2* exon 2 (Figure 1A) following our previous approach described for generation of zebrafish proneural Cre drivers in *ascl1b*, *olig2*, and *neurod1* ^3^. The knock-in cassette contained a linked secondary marker for lens-specific expression of a fluorescent reporter to facilitate allele identification by fluorescence screening on a stereomicroscope (Figure 1B). The *pPRISM-2A-cre, gcry1:mRFP* and *pPRISM-2A-creERT2, gcry1:eGFP* GeneWeld donor plasmids are part of the GeneWeld pPRISM (plasmids for Precise Integration with Secondary Markers) vector suite available at AddGene/Jeffrey Essner Lab Materials.

**Figure 1.**
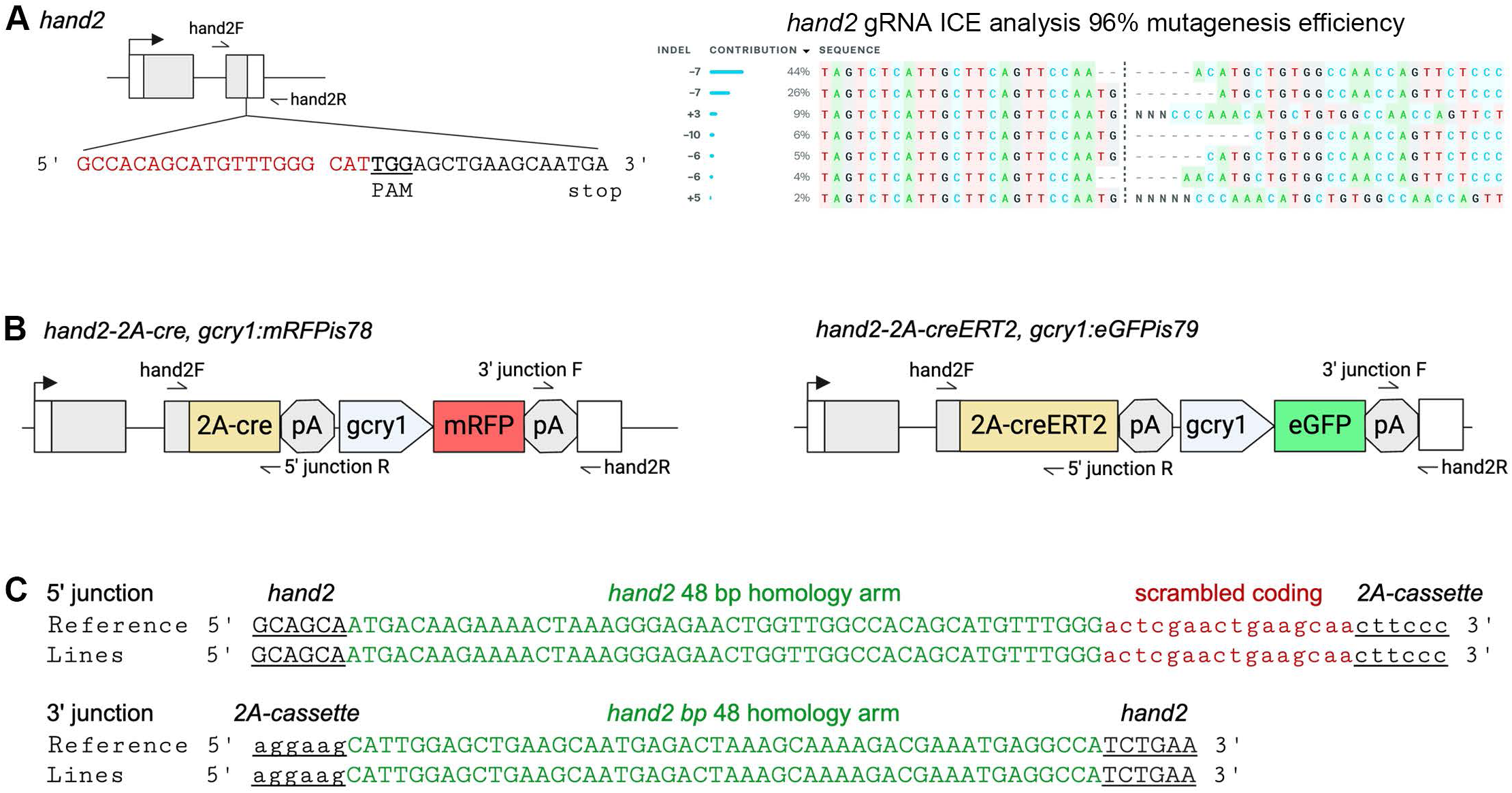
*hand2* 2A-cre and 2A-creERT2 knock-in lines generated with GeneWeld CRISPR/Cas9 targeted integration. **A** *hand2* diagram showing sequence of exon 2 gRNA in red relative to the *hand2* translation stop codon TGA. Space, Cas9 cut site. PAM in bold and underlined. Primers for generating PCR amplicon to test *hand2* gRNA mutagenesis efficiency are labeled hand2F and hand2R. Results of ICE analysis of Sanger sequenced PCR amplicon from showing 96% mutagenesis efficiency and range of indel mutations at the *hand2* gRNA target site. **B** Diagrams of recovered *hand2-2A-cre, gcry1:mRFPis78* and *hand2-2A-creERT2, gcry1:eGFPis79* lines with primers used for 5’ and 3’ junction PCR analysis. **C** Sanger sequence results of 5’ and 3’ genomic DNA - cassette knock-in junction PCR amplicons from Founders and F2 adults. 48 bp homology arms are shown in green. 5‘ homology arm contained an additional 17 nucleotides to complete the *hand2* coding sequence, scrambled, shown in red.

Zebrafish *hand2* was targeted with a forward strand gRNA 5’- *GCCACAGCATGTTTGGGCAT**<u>TGG</u>***-3’ (PAM underlined and in bold) that cuts 17 bp upstream of the *hand2 TGA* translation stop codon and shows 96% mutagenesis efficiency in individual injected WIK embryos (Figure 1 A). The *2A-cre* and *2A-creERT2* donor plasmids contained a 69 bp 5’ homology arm with scrambled codons, to prevent cutting by the genomic gRNA, followed by the remainder of the *hand2* coding sequence, and a 48 bp 3’ homology arm. 8 embryos expressing the lens-expressing fluorescent secondary marker from each outcrossed adult founder were tested for precise integration alleles by PCR junction analysis and Sanger sequencing following established protocols (Wierson 2020; Welker 2021; Almeida 2021). 23% (10/43) of *hand2-2A-cre, gcry1:mRFP* and 11% (5/46) of *hand2-2A-creERT2, gcry1:eGFP* adults transmitted a germline allele with expression of the lens-specific secondary marker. 20% of positive adults transmitted a precise integration allele: 2/10 *hand2-2A-cre, gcry1:mRFP* adults and 1/5 *hand2-2A-creERT2, gcry1:eGFP* adults. This corresponds to an overall frequency of 5% (2/43) and 2% (1/46) of all adults screened, respectively (Table 1). Individual F1 and F2 adults harboring precise integration alleles were used to establish lines *hand2-2A-cre, gcry1:mRFP^is^*^78^ and *hand2-2A-creERT2, gcry1:eGFP^is^*^79^. PCR on F2 adults with primers flanking *hand2* genomic DNA-cassette junctions (Figure 1 B) generated amplicons that showed the expected sequences corresponding to precise integration at both 5’ and 3’ sides of the integration (Figure 1 C) in both lines. To confirm on-target integration, Whole Genome Sequencing was performed using PacBio long-read sequencing on genomic DNA isolated from *hand2-2A-cre, gcry1:mRFP^is^*^78*/+*^ and *hand2-2A-creERT2, gcry1:eGFP^is^*^79*/+*^ F3 adults. Mapping the *hand2-2A-cre* and *hand2-2A-creERT2* vector sequences revealed a single integration of the *2A-cre/creERT2*, secondary marker cassette at the *hand2* 3’ target site in *hand2-2A-cre, gcry1:mRFP^is^*^78*/+*^ and *hand2-2A-creERT2, gcry1:eGFP^is^*^79*/+*^. *hand2-2A-cre, gcry1:mRFP^is^*^78^ and *hand2-2A-creERT2, gcry1:eGFP^is^*^79^ heterozygous and homozygous larva developed normally through larval stages without morphological defects or developmental delay, and homozygotes survive to adulthood. For simplicity the lines are referred to as *hand2-2A-cre* and *hand2-2A-creERT2*.

**Table 1.**
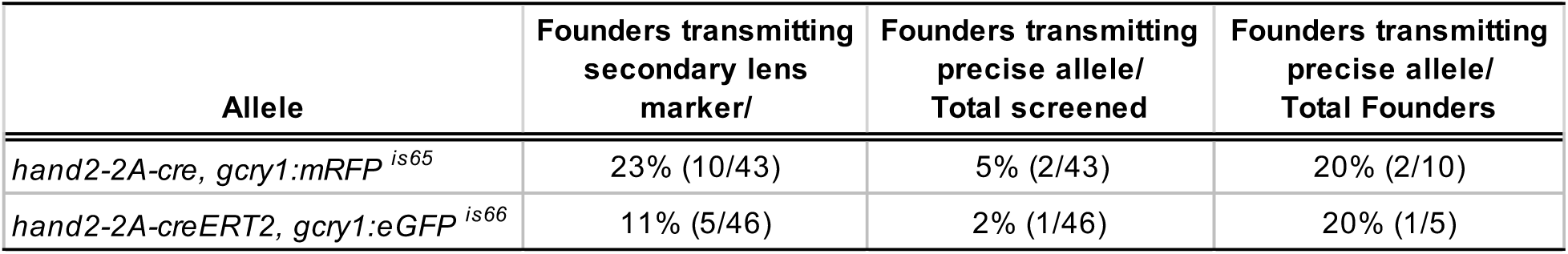
Recovery of *hand2-2a-cre* and *hand2-2A-creERT2* alleles by GeneWeld CRISPR precision targeted integration. From all Founder adults that transmit a germline allele expressing the lens-specific secondary marker, 20% transmitted precise integration alleles where both 5’ and 3’ junctions show seamless, scarless integration.

### *hand2-2A-cre* recombinase lineage tracing reveals mesoderm and ectoderm derived lineages in the embryo

To assess *hand2-2A-cre* lineage specific Cre activity the line was crossed with the ubiquitously expressed floxed switch reporter *Tg(3.5ubi:loxP-lacZ-loxP-eGFP)^cn^*^2^ ^25^, referred to as *ubi:Hulk*. Cre recombinase activity results in excision of the floxed STOP cassette and ubiquitous GFP expression ^25^. At 22 hpf, in *hand2-2A-cre*; *ubi:Hulk* embryos GFP expression was detected in mesoderm-derived tissues, including the heart and migrating mesothelial progenitors (Figure 2 A, A’ large arrow; Figure 2 B, B’ large arrow) and the fin buds (Figure 2 B, B’ arrowheads). GFP signal was observed in a layer surrounding the yolk extension (Figure 2 A, A’ small arrowheads) indicative of mesothelium as previously described ^18,26^. GFP could also be detected in cells dorsal to the yolk periderm where the lateral plate mesoderm flanks the tissue that will become the primitive gut ^17^ (Figure 2 A, A’ small arrows). GFP positive cells were not detected in the gut primordium that lies dorsal to the yolk and below the vascular cell layers (Figure 2 C, C’ arrows).

**Figure 2.**
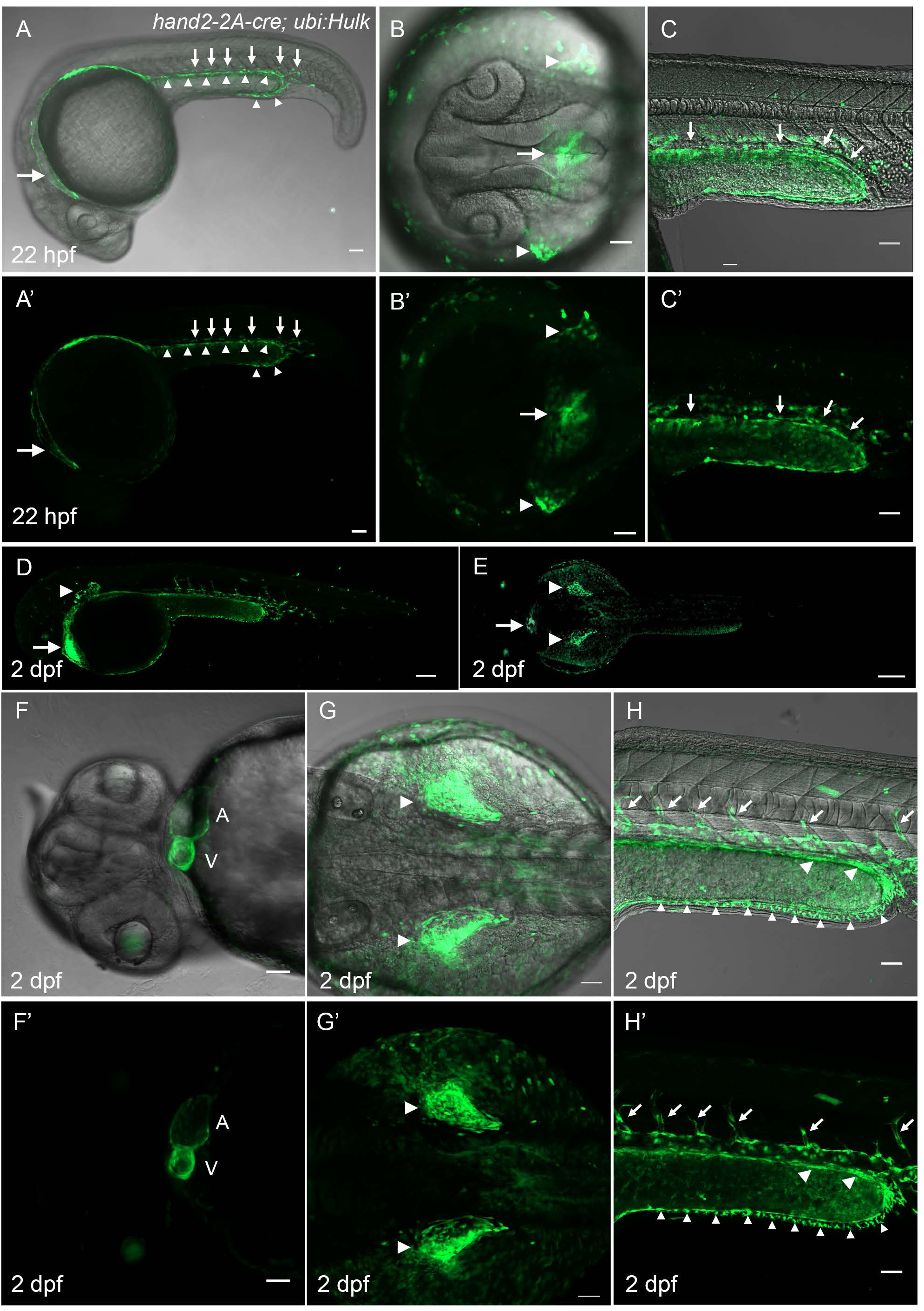
*hand2* lineage tracing live confocal imaging at 22 hpf and 2 dpf in *hand2-2A-cre, gcry1:mRFP; ubi:Hulk-lox-Stop-Lox-eGFP* embryos. **A, A’** GFP expression in 22 hpf embryo in mesothelium (large arrow), mesoderm-derived vessels (small arrows), and yolk periderm (small arrowheads). **B, B’** Higher magnification dorsal view at 22 hpf shows GFP expression in the heart (arrows) and fin buds (arrowheads). **C, C’** Higher magnification lateral view of posterior yolk extension show GFP is absent from cell layers of the presumptive gut (small arrows). **D** Lateral and **E** dorsal views of 2 dpf embryos show strong GFP expression in the heart (arrows) and fin buds (arrowheads). **F, F’** Ventral view of the heart at 2 dpf shows GFP in the atrium (A) and ventricle (V). **G, G’** Higher magnification dorsal views of GFP in the fin buds (arrowheads). **H, H’** Higher magnification lateral views of 2 dpf mid trunk region show GFP in sprouting vessels from the posterior cardinal vein (small arrows), in presumptive cell layer of the developing gut (arrowheads) and pectoral fin fold (small arrowheads). Scale bars 50 μm.

At 2 days post-fertilization (dpf), GFP was detected in the heart, the pericardium surrounding the heart (Figure 5 D, E, arrows), and in both atrial and ventricular chambers (Figure 2 F, F’, labels A and V). Strong GFP expression was detected in the pectoral fin buds (Figure 2 D, E, G, G’ arrowheads) and in the gut (Figure 2 H, H’ arrowheads). GFP was also present on the ventral side of the yolk extension in the pre-anal fin fold (PAFF) in the PAFF lateral plate mesoderm-derived fibroblasts ^26^ (Figure 2 H, H’ small arrowheads) as previously reported in Tzung et al., 2023 ^26^. Notably, GFP expression was present in the posterior cardinal vein (PCV) and in the intersegmental vessels sprouting from the PCV (Figure 2 H, H’ arrows).

Each of these patterns lends support to previous studies demonstrating a requirement for *hand2* in the myocardium, aortic arch, branchial arches, forelimb/pectoral fin mesenchyme, PAFF fibroblasts, smooth muscle and gut looping, sympathetic neurons, and the posterior vasculature ^9,11,12,14–16,26^. Overall, analysis of *hand2-2A-cre* lineage tracing indicates *hand2* progenitors originate in the lateral plate mesoderm and ectoderm and contribute to mesodermal-derived tissues in multiple organs during embryogenesis.

In 3 dpf larva, GFP persisted in presumptive mesoderm derived tissues in the heart, pectoral fins (Figure 3 A arrow, arrowhead), posterior cardinal vein and intersegmental vessels (Figure 3 A, B small arrows), and surrounding the gut (Figure 3 B small arrowheads). GFP was present in both the heart atrial and ventricular myocardium (Figure 3 C, C’, labels A and V), the branchial arches (Figure 3 C, C’ label PA), the pectoral fins (Figure 3 D, D’ large arrowheads), and the anterior portion of the digestive tube (Figure 3 D’, small arrowheads). At 5 dpf, strong GFP expression persisted in the branchial arches (Figure 3 E, F label PA) and again in the myocardium in the atrium and ventricle (Figure 3 E label V; Figure F labels A, V). High levels of GFP were detected in the bulbus arteriosus at the outflow tract (Figure 3 G, G’ label BA). A low level of GFP was present in the atrioventricular valve (Figure 3 G, G’ arrowhead), possibly due to cell density in the heart tissue. In the 6 dpf liver (Figure 3 H, H’ dashed area), GFP-positive cells with morphology akin to hepatic stellate cells were present (Figure 3 H, H’ asterisks).

**Figure 3.**
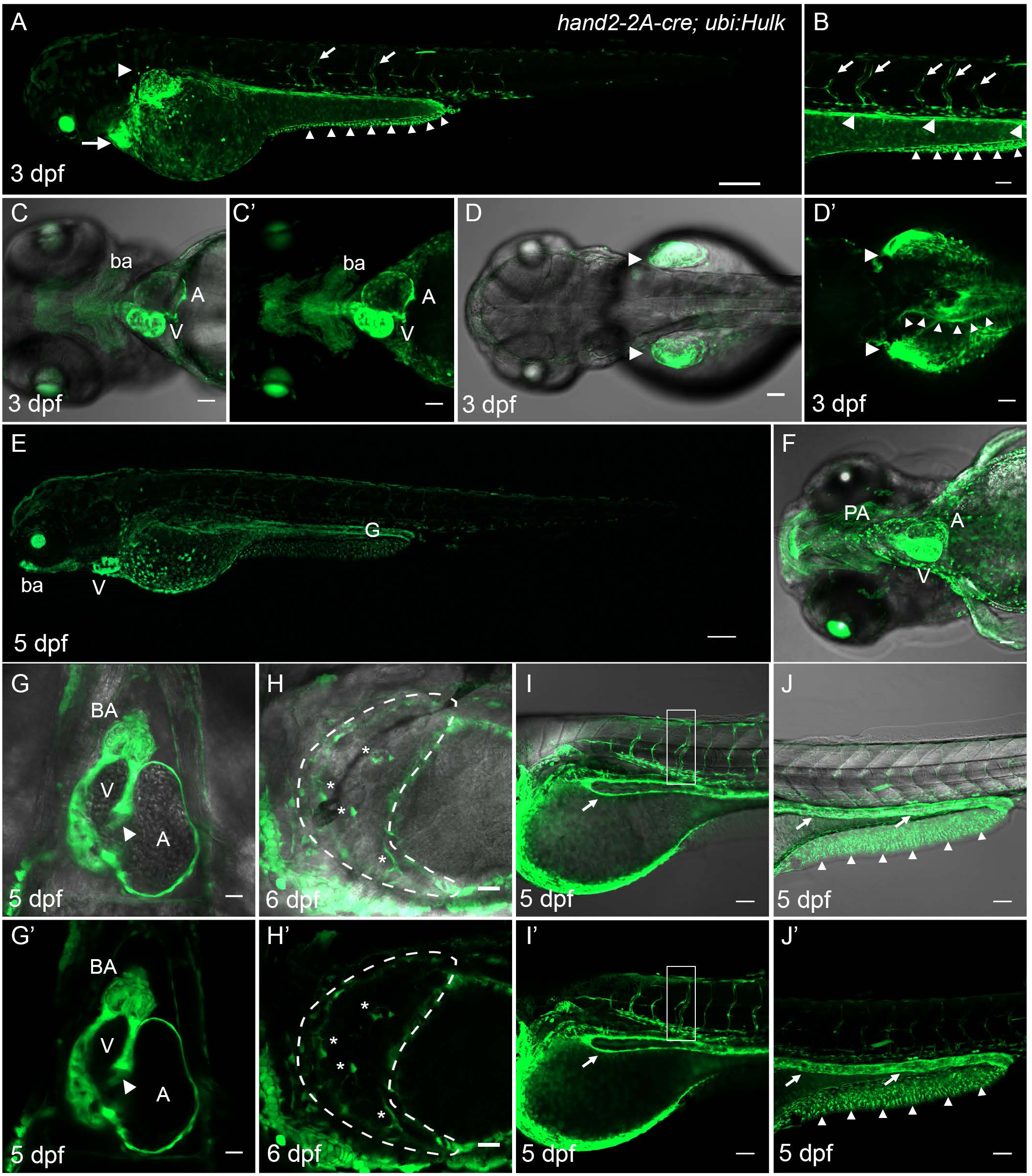
*hand2* lineage tracing in 3 and 5 dpf *hand2-2A-cre, gcry1:mRFP; ubi:Hulk-lox-Stop-Lox-eGFP* larvae. **A-D** Live confocal imaging of GFP expression in 3 dpf *hand2-2a-cre; ubi:Hulk* larvae. **A** Strong GFP expression is detected in heart (arrow), fin (arrowhead) and vasculature (small arrows), and pectoral fin fold (small arrowheads). **B** Higher magnification of the trunk shows GFP expression in intersegmental vessels and posterior cardinal vein (arrows), the developing intestine (arrowheads), and pectoral fin fold (small arrowheads). **C, C’** Ventral view of GFP expression in the branchial arches (ba) and heart ventricle (V) and atrium (A) chambers. **D** Dorsal view of GFP expression in the pectoral fins (arrowheads). **D’** More dorsal focal plane showing GFP expression in pectoral fins (large arrowheads) and in the developing gut (small arrowheads). **E – J** Live confocal imaging of GFP expression in 5 dpf *hand2-2a-cre; ubi:Hulk* larvae. **E** Strong GFP expression in the branchial arches (ba), heart ventricle (V) and along the lining of the developing gut (G). **F** Ventral view shows GFP expression in the branchial arches (PA) and strong expression in the heart ventricle (V) and atrium (A). **G, G’** GFP expression is present in the heart bulbus arteriosus (BA), ventricle (V), atrium (A), and atrioventricular valve (arrowhead). **H, H’** GFP expression in mesothelial tissue surrounding the liver (dashed line) and in hepatic stellate cells (asterisks). **I, I’** GFP expression in the vasculature in the anterior trunk in intersegmental vessels and posterior cardinal vein (boxed region) and in the lining of the anterior gut (arrow). **J, J’** Strong GFP expression extending along the length of the intestine (small arrows) and in the pectoral fin fold (small arrowheads). Scale bars: A, B, C, C’, D, D’ E, F, 50 μm; G, G’, H, H’, I, I’, J, J’, 20 μm.

Vascular GFP expression was detected in intersegmental vessels and the posterior cardinal vein (Figure 3 I, I’ boxed region). GFP labeling persisted along the length of the intestine (Figure 3 I, I’, J, J’ small arrows) and in the PAFF fibroblasts (Figure 3 J, J’ small arrowheads).

To more closely examine *hand2* lineage tracing in the developing gut and PCV, transverse sections of fixed 24 hpf, 48 hpf, and 72hpf *hand2-2A-cre, gcry1:nlsmRFP*; *ubi:lox-GFP-lox-mCherry* (*ubi:Switch*) embryos and larvae were labeled with DAPI and Phalloidin (Figure 4). In the 24 hpf embryo mCherry was present in the Lateral Plate Mesoderm flanking the region where the endoderm-derived primitive gut will form (Figure 4 A label LPM). A few mCherry positive cells were present in the neural tube (Figure 4 A Neural tube), which may represent hand2 neural crest cells. At 48 hpf mCherry was present in the LPM surrounding the primitive gut (Figure 4 B, label LPM, g). At this stage mCherry could also be detected in the PCV (Figure 4 B, PCV), consistent with live imaging of GFP using the *ubi:Hulk* Switch line (Figure 3). Transverse sections located at anterior, mid, and posterior positions along the trunk of a 72 hpf larva showed mCherry expression in the smooth muscle layer surrounding the intestine (Figure 4 C, D, E Intestine). mCherry was also present in cells in the liver (Figure 4 C, liver), which may represent hepatic stellate cells as previously reported ^27^, and in the PCV (Figure 4 D, E PCV).

**Figure 4.**
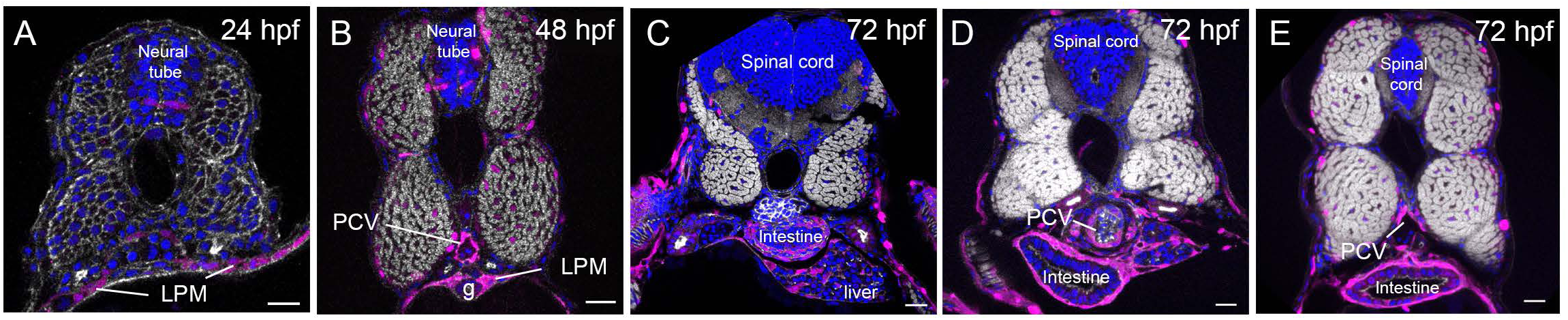
Vibratome transverse sections through the trunk of 24 hpf, 48 hpf and 72 hpf *hand2-2A-cre, gcry1:mRFP; ubi:Switch lox-GFP-lox-mCherry* embryos and larvae. **A – E** 150 μm transverse sections labeled with DAPI (blue) and Phalloidin (White). **A** 24 hpf embryo shows mCherry (magenta) expression in the Lateral Plate Mesoderm (LPM) flanking the endoderm that will form the primitive gut. mCherry is present in a small number of cells in the neural tube. **B** 48 hpf embryo shows mCherry expression in the LPM surrounding the primitive gut (g). mCherry expression is present in the Posterior Cardinal Vein (PCV) and in a few cells in the neural tube. Nuclear mCherry present in the muscle may represent ectopic expression of the *gcry1:nls-mRFP* linked secondary marker in the *hand2-2A-Cre* knock-in allele. **C – E** Transverse sections along the anterior-posterior axis of a 72 hpf larva. **C** Anterior transverse section shows mCherry in the mesoderm-derived smooth muscle surrounding the Intestine, and in cells within the liver**. D** Mid trunk transverse section, and **E** Posterior trunk transverse section, shows mCherry expression in the smooth muscle surrounding the Intestine and in the wall of the PCV. n=8 embryos or larva per time point. Scale bars A, B, D, E 20 μm; C 30 μm.

Together, *hand2-2A-cre* lineage tracing through 5 dpf was consistent with *hand2*-positive lateral plate mesoderm progenitors contributing to the branchial arches, heart, and pectoral fin ^11,27^, epithelial lateral plate mesoderm ^14^ and intestinal smooth muscle ^17^, mesothelium ^18^, the liver ^27^, and the PAFF ^26^. In addition to the expected patterns of *hand2-2a-cre* lineages, our analyses confirmed *hand2*-expressing progenitors contribute to the posterior venous vasculature.

### *hand2-2A-creERT2* allows Tamoxifen-regulated lineage tracing that recapitulates *hand2-2A-cre* patterns

To demonstrate the *hand2-2A-creERT2* driver could be used for 4-OH-tamoxifen (4- OHT)-controlled temporal induction of *hand2* lineage tracing, CreERT2 recombinase activity was induced at the embryonic shield stage to test for recapitulation of lineages labeled by *hand2-2A-cre*. The *hand2-2A-creERT2, gcry1:eGFP* driver was crossed to the ubiquitous switch reporter *Tg(-3.5ubb:loxP-EGFP-loxP-mCherry)cz1701*^28^ (*ubi:Switch*), which switches from GFP to mCherry expression after recombination excision of a floxed GFP cassette ^28^. Embryos were placed in 10 μm 4-OHT in embryo medium at shield stage and *hand2-2A-creERT2; ubi:Switch* embryos were imaged at 24 hpf and 2 dpf (Figure 5). *hand2-2A-creERT2; ubi:Switch* 24 hpf embryos showed mCherry expression in the developing heart (Figure 5 A, A’ large arrow; Figure 5 B, B’ arrow), mesothelium and PAFF (Figure 5 A, A’ arrowheads), and developing posterior vasculature (Figure 5 A, A’ small arrows) as observed with *hand2-2A-cre, ubi:Hulk* embryos (Figure 2). Higher magnification of the lateral trunk shows mCherry is present in the migrating PAFF fibroblasts and developing vasculature located below and above the cell layers of the primitive gut, which remain positive for GFP as unrecombined cells (Figure 5 C, C’ arrows).

**Figure 5.**
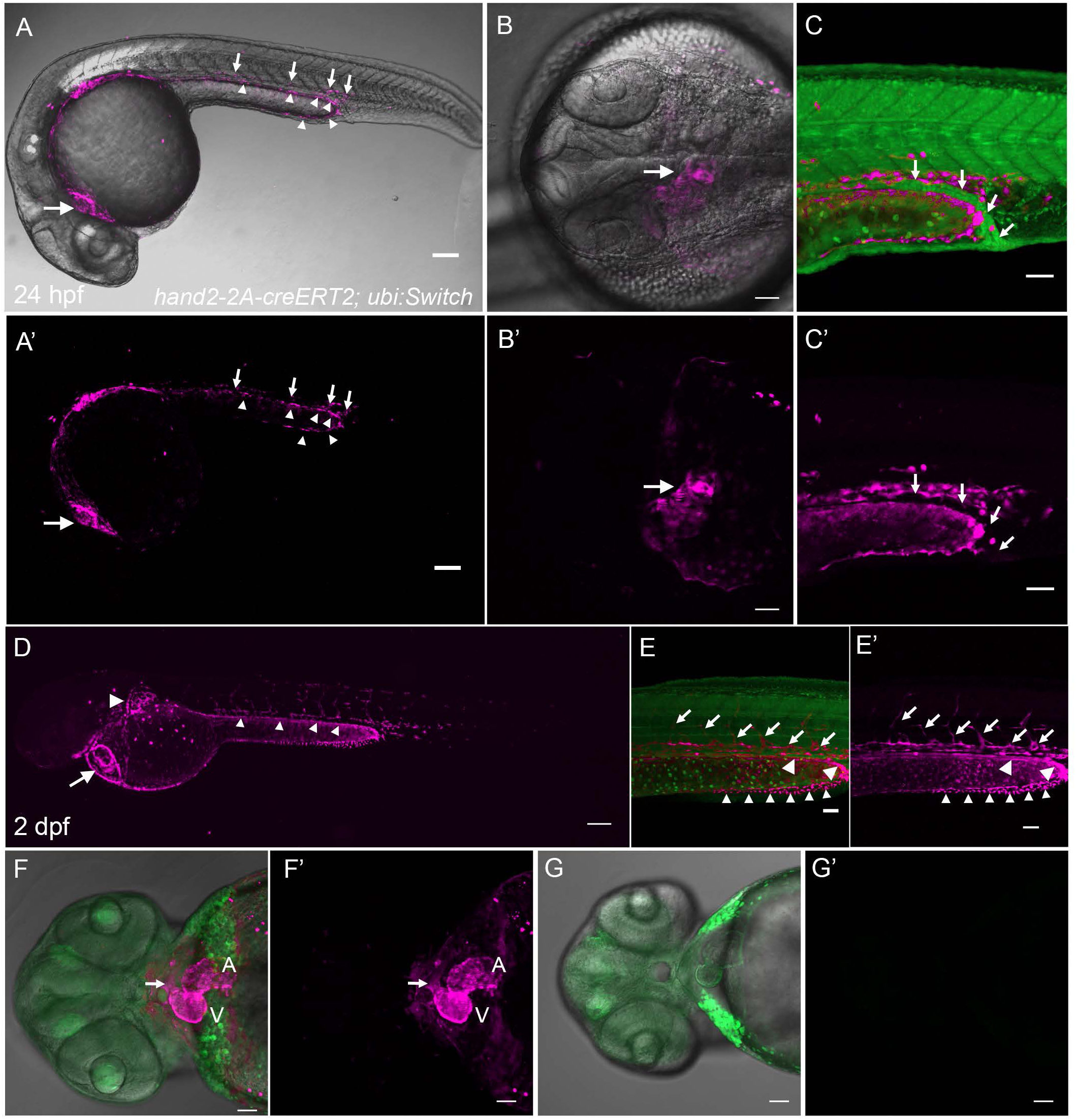
*hand2-2A-creERT2, gcry1:eGFP; ubi:Switch-lox-GFP-STOP-lox-mCherry* lineage tracing in 24 hpf and 2 dpf embryos after tamoxifen regulated switch at 6 hpf shield stage. **A – C** Live confocal imaging of *hand2-2A-creERT2* mCherry (magenta) labeled lineages in 24 hpf embryos. **A, A’** Lateral view showing mCherry expression in the heart (large arrow), vasculature (small arrows) and fibroblasts of the pre-anal fin fold (PAFF) (small arrowheads). **B, B’** Dorsal view showing mCherry expression in the developing heart (arrow). **C, C’** Absence of GFP to mCherry switching in the cell layers of the primitive gut (arrows). **D – F** Live confocal imaging of *hand2-2A-creERT2* mCherry labeled lineages in 2 dpf embryos. **D** Lateral view showing mCherry expression in the pericardium and heart (arrow), fin bud (large arrowhead) and PAFF (small arrowheads). **E, E’** Higher magnification lateral view of trunk shows mCherry in posterior cardinal vein and intersegmental vessels (arrows), the primitive gut (arrowheads), and the PAFF (small arrowheads). **F, F’** Ventral view embryo shows mCherry expression in the atrium (A) and ventricle (V) of the heart and a low level of expression in the presumptive heart outflow tract (arrow). **G, G’** mCherry expression is not detected in 2 dpf control *hand2-2A- creERT2; ubi:Switch* embryo treated with EtOH vehicle alone. Scale bars: A, A’ 100 μm; B, B’, C, C’, D, D’ E, F, F’, G, G’ 50 μm.

At 2 dpf *hand2-2A-creERT2; ubi:Switch* embryos showed mCherry expression in the heart and pericardium surrounding the heart (Figure 5 D arrow), the fin bud (Figure 5 D large arrowhead), and in the gut (Figure 5 D small arrowheads). mCherry was present in the posterior cardinal vein and sprouting intersegmental vessels (Figure 5 E, E’ small arrows), the primitive gut (Figure 5 E, E’ arrowheads), and the PAFF (Figure 5 E, E’ small arrowheads). In the heart mCherry was present in both atrium and ventricle (Figure 5 F, F’ labels A, V) and detected at a low level in the outflow tract region of the heart (Figure 5 F, F’ small arrow). In the absence of 4- OHT treatment, mCherry expression was not detected in any tissue in the developing embryo (Figure 5 G, G’), demonstrating switching and induction of mCherry expression was dependent on 4-OHT induction of Cre recombinase activity.

The robustness of *hand2-2A-creERT2* for 4-OHT-regulated switching activity was further tested with the heat shock-based Switch reporter *line hsp70l:lox-STOP-lox-GFP, cryaa:Venus*^29^. Cre recombinase activity will lead to excision of a floxed STOP cassette, however, GFP expression is only induced after activation of the *hsp70l* promoter by heat shock treatment ^29,30^, providing a second layer of inducible lineage tracing. *hand2-2A-cre; hsp70l:Switch* embryos treated with 4-OHT at shield stage were heat shocked at 37°C for 1 hour beginning 3 hours before imaging, as previously performed ^30^. GFP lineage labeling after heat shock at 1.5 dpf and 2.5 dpf showed GFP in the heart (Figure 6 A, A’, B, B’, C, C’ large arrows), pectoral fin bud (Figure A, A’, C, C’ large arrowheads), branchial arches (Figure 6 B, B’ large arrowheads), posterior cardinal vein (Figure 6 A, A’, C, C’ small arrows), and mesothelium and PAFF (Figure 6 A, A’, C, C’ asterisks). Induction of *hsp70l:Switch* GFP reporter expression also revealed lineage tracing in structures that arise in the late-stage embryo. In 2.5 dpf larva GFP was detected in PCV-derived intersegmental vessels (Figure 6 C, C’ small yellow arrows), PAFF fibroblasts (Figure 6 C, C’ large yellow arrows), and the developing gut (Figure 6 C, C’ small arrowheads). GFP was not detected in 1.5 hpf or 2.5 dpf embryos in the absence of 4-OHT treatment, demonstrating switching by CreERT2 recombination was necessary for induction of GFP expression. Together, these results demonstrate combining 4-OHT regulation of CreERT2 activity with temporal induction of our heat shock *hsp70l*-based reporter will allow lineage tracing in structures that form after a defined stage of development.

**Figure 6.**
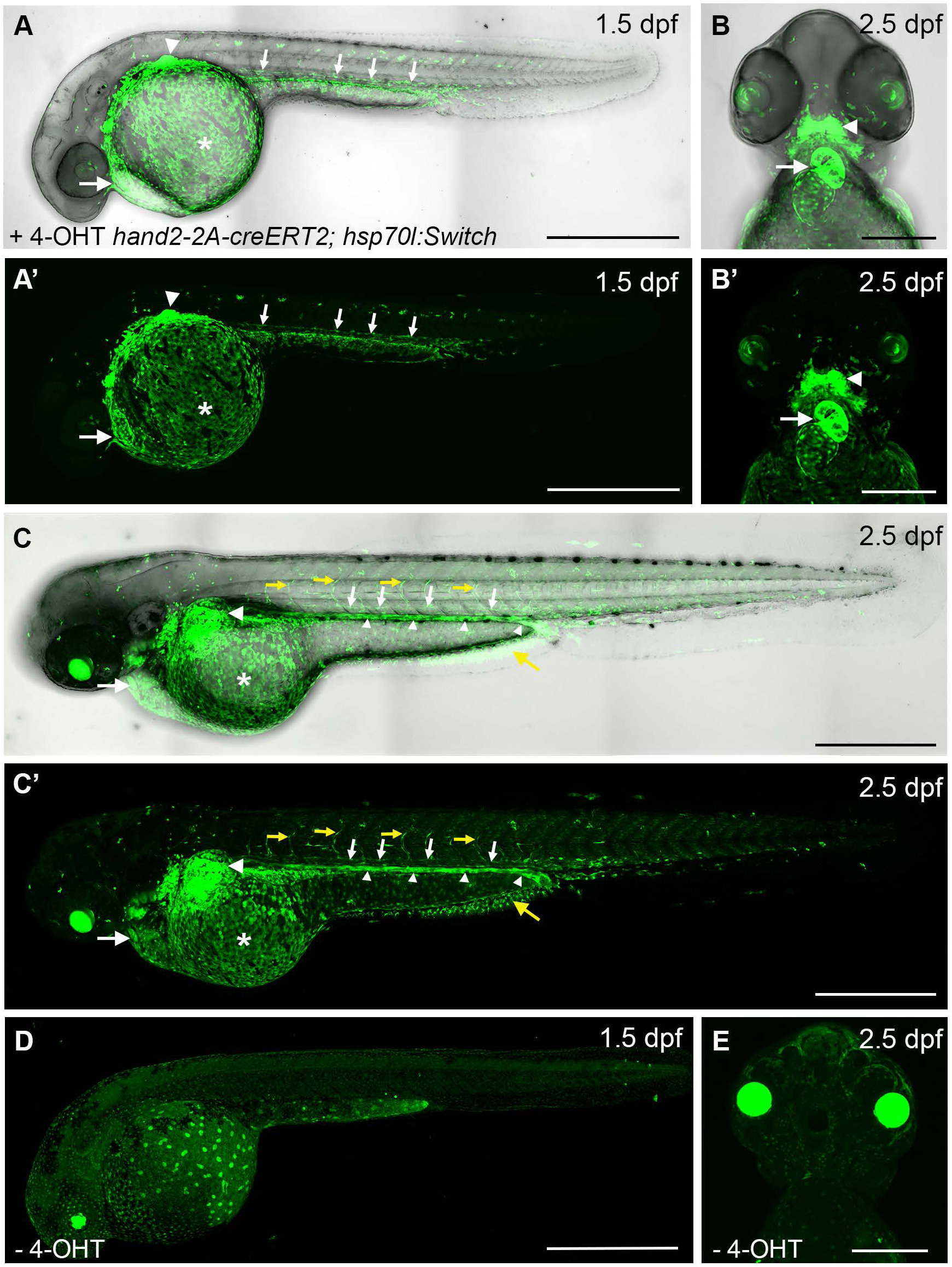
Tamoxifen and heat shock regulated lineage tracing in *hand2-2A-creERT2; hsp70l:lox-STOP-lox-GFP* embryos at 1.5 and 2.5 dpf. **A, A’** Live confocal imaging of *hand2-2A-creERT2; hsp70l:lox-STOP-lox-GFP* embryos after 4-OHT treatment at shield stage and heat shock at 1.5 dpf. GFP expression is present in the heart (large arrow), pectoral fin (large arrowhead), Posterior Cardinal Vein (small arrows), and PAFF (asterisks). **B, B’, C, C’** Live confocal imaging of *hand2-2A-creERT2; hsp70l:lox-STOP-lox-GFP* embryos after 4-OHT treatment at shield stage and heat shock at 2.5 dpf. **B, B’** Ventral view of 2.5 dpf embryo showing GFP expression in the cardiac atrium and ventricle (arrows) and branchial arches (arrowheads). **C, C’** Lateral view of 2.5 dpf embryo showing GFP expression in the heart (large arrows), pectoral fin bud (large arrowheads), Posterior Cardinal Vein (small arrows), venous intersegmental vessels (small yellow arrows), yolk periderm (asterisks), PAFF (large yellow arrows), and primitive gut (small arrowheads). **D, E** Live confocal imaging of 1.5 dpf (**D**) and 2.5 dpf (**E**) *hand2-2A-creERT2; hsp70l:lox-STOP-lox-GFP* control embryos that were not treated with 4-OHT. Scale bars: A, A’, C, C’, D 500 μm; B, B’, E 200 μm.

### *hand2-2A-creERT2* Tamoxifen-regulated lineage tracing of embryonic vs. larval hand2 progenitors

To determine if *hand2* expression persists in select progenitors in post-embryonic organs and contributes to larval organogenesis, lineage tracing at 5 dpf was compared after 4- OHT treatment at shield stage vs. treatment in 3 dpf larva. 4-OHT induced switching of *hand2-2A-creERT2; ubi:Switch* during shield stage resulted in lineage tracing at 5 dpf that matched the complete pattern of *hand2-2A-cre* (Figure 7 A – E). Strong mCherry signal could be detected in the brachial arches (Figure 7 A, A’ label PA), heart (Figure 7 A, A’ arrow), pectoral fins (Figure 7 A, A’ arrowheads), bulbus arteriosus (Figure 7 B, B’ label BA), myocardium of the atrium and ventricle (Figure 7 B, B’ label A, V) and atrioventricular valve (Figure 7 B, B’ arrowhead). mCherry could also be detected in the presumptive pericardium (Figure 7 B, B’ small arrows). At 6 dpf mCherry was detected in liver cells with hepatic stellate cell morphology (Figure 7 C, C’ asterisks) and cells in the mesothelium surrounding the liver (Figure 7 C, C’ dashed shapes). mCherry expression was once more present in the posterior cardinal vein and intersegmental vessels (Figure 7 D, D’, E, E’ small arrows). In the developing gut mCherry could be detected in the intestinal bulb (Figure 7 D, D’ large arrow), and the middle and distal intestine (Figure 7 D, D’, E, E’ large arrowheads), which presumably corresponds to intestinal smooth muscle as shown in transverse cross sections in 3 dpf hand2-2A-cre; ubi:Switch larva (Figure 4). mCherry was highly expressed in the PAFF (Figure 7 D, D’, E, E’ asterisks).

**Figure 7.**
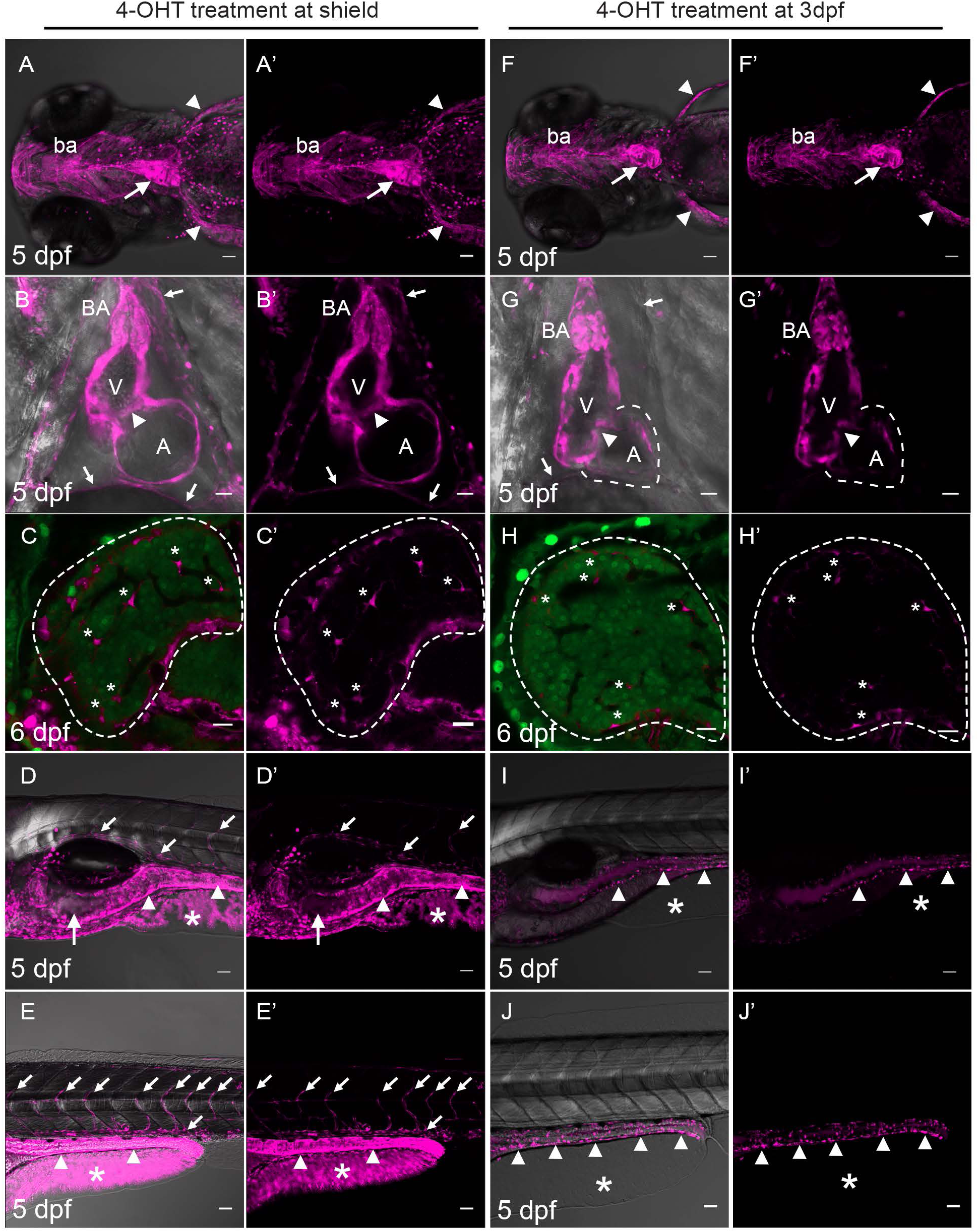
Comparison of *hand2-2A-creERT2* lineage tracing in 5dpf larvae after induction at shield stage vs. 3 dpf. **A –E** Live confocal imaging of 5 dpf *hand2-2A-creERT2; ubi:Switch* embryos treated with 4-OHT at 6 hpf. **A, A’** Ventral view showing mCherry (magenta) expression in branchial arches (ba), heart (arrow) and pectoral fins (arrowheads). **B, B’** mCherry expression in the cardiac bulbus arteriosus (BA), ventricle (V), atrium (A), atrioventricular valve (arrowhead), and pericardium (small arrows). **C, C’** mCherry expression in the mesothelium surrounding the liver (dashed line) and hepatic stellate cells (asterisks). **D, D’** Anterior trunk region showing mCherry expression in the intersegmental vessels and posterior cardinal vein (small arrows), intestinal bulb (arrow), mid intestine (arrowheads), and PAFF fibroblasts (asterisk). **E, E’** Mid-trunk region showing mCherry expression in the intersegmental vessels and posterior cardinal vein (small arrows), intestine (arrowhead), and PAFF (asterisk). **F – J** Live confocal imaging of 5 dpf *hand2-2A-creERT2; ubi:Switch* embryos treated with 4-OHT at 3 dpf. **F, F’** Ventral view showing mCherry expression in branchial arches (PA), heart (arrow) and pectoral fins (arrowheads). **G, G’** mCherry expression in the heart bulbus arteriosus (BA), ventricle (V), atrioventricular valve (arrowhead), and atrium (A, dashed outline), absent from the pericardium (small arrows). **H, H’** mCherry expression in liver hepatic stellate cells (asterisks). Dashed line outlines liver. **I, I’** Anterior and (**J, J’**) mid trunk regions showing mCherry expression present in gut enteric neurons (arrowheads) and absent from the PAFF (asterisk). Scale bars: B, B’, E, E’. F, F’, G, G’ J, J’, K, K’, 50 μm; C, C’. D, D’, H, H’, I, I’, 20 μm.

In contrast to the broad labeling after induction of Cre recombinase at shield stage, treatment with 4-OHT at 3 dpf resulted in mCherry expression in a subset of larval organs (Figure 7 F - J). mCherry expression was detected in the branshial arches, heart and pectoral fins (Figure 7 F, F’ label PA, arrow, and arrowheads), indicating *hand2* continues to be expressed in cardiac muscle and mesenchymal tissue. Expression in the branchial arches was less extensive in comparison to switching at shield stage and appeared limited to the outer edges of the arches. Within the heart, mCherry signal was strong in the bulbus arteriosus and ventricle (Figure 7 G, G’ label BA, V) and present in the atrioventricular valve (Figure 7 G, G’ arrowhead). In contrast to lineage tracing after switching at shield stage, mCherry was only detected in a few cells of the atrium in the heart (Figure 7 G, G’ label A, dashed outline) and was absent from the presumptive pericardium (Figure 7 G small arrows). In the liver, mCherry expression was present in hepatic stellate cells (Figure 7 H, H’ asterisks) but absent from cells mesothelial cells surrounding the liver. These results indicate *hand2* lineages that contribute to mesothelial tissues are derived from early progenitors in the lateral plate mesoderm.

Structures that were labeled by early *hand2* progenitors, but did not show mCherry expression after induction in the 3 dpf larva, included the anterior abdomen, presumed mesodermal tissue surrounding the gut, and the PAFF (Figure 7 I, I’ asterisk). Expression was detected in cells surrounding the gut, extending from the middle to the distal end of the intestine, with morphology reminiscent of enteric neurons (Figure 7 I, I’, J, J’ arrowheads). Notably, mCherry was entirely absent from the posterior venous vasculature (Figure 7 I, I’, J, J’). These results indicate *hand2* progenitors that persist in post embryonic tissues may be restricted to pharyngeal arches, heart, and liver mesodermal tissues, and the intestine, consistent with previous *in situ* hybridization analysis of hand2 expression in 3 dpf ^26^ and 4 dpf ^27^ zebrafish larva.

To identify cell types derived from late *hand2*-expressing progenitors contributing to the larval liver and gut, confocal imaging was performed on tissues from 5 dpf fixed larva (Figure 8). mCherry-positive cells with the morphology of hepatic stellate cells were present in the liver (Figure 8 A, A’ arrows), consistent with a previous report of *hand2* labeling of hepatic stellate cells ^27^, and a small number of mCherry-positive mesothelial-like cells were detected at the surface. mCherry labeled cells with the morphology of enteric neurons and neurite extensions were present encircling the gut (Figure 8 B, B’), indicative of enteric neurons derived from the vagal neural crest ^31^. These results were consistent with the observation that late *hand2* progenitors in the larva reside in mesodermal and enteric neural tissue.

**Figure 8.**
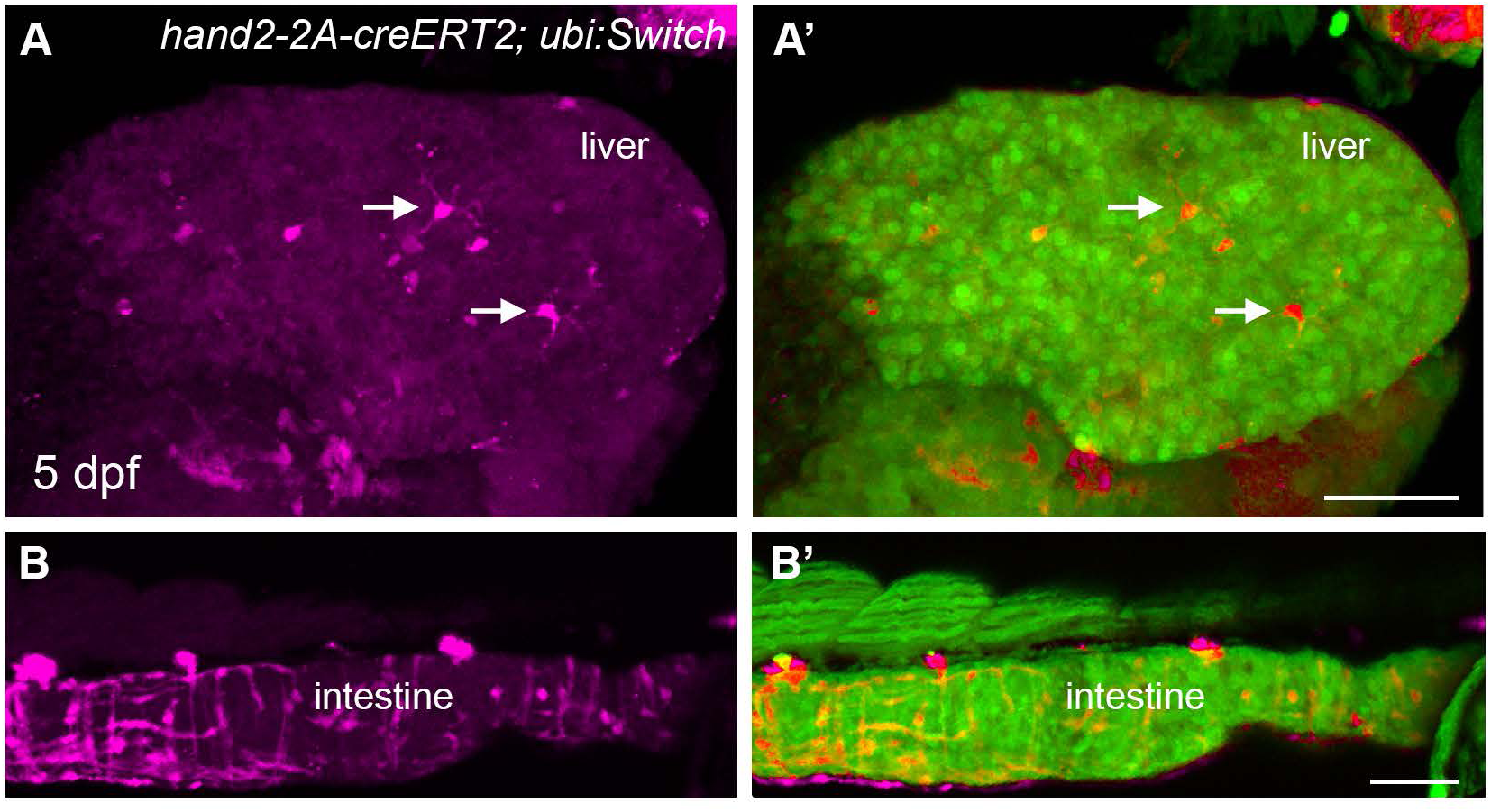
mCherry expression in *hand2-2A-creERT2; ubi:Switch* fixed 5 dpf larval liver and intestine after induction at 3 dpf. **A, A’** Confocal 3D projection of z-stack sections of mCherry (magenta) expression in liver hepatic stellate cells (arrows). **B, B’** Confocal 3D projection of z-stack sections of mCherry expression in enteric neuron cell bodies and neurite extensions surrounding the intestine. Scale bars: A, A’, B, B’ 50 μm.

### *hand2* lineage contributes to the posterior cardinal vein and venous-derived intestinal vasculature

To better define the temporal and spatial contribution of *hand2*-derived progenitors to the different vasculature beds, we crossed *hand2:CreERT2* to the ubiquitous *actb2:loxP-BFP-loxP-dsRed* switch line carrying the endothelial reporter *fli1:GFP* in the background. We first added 4- OHT at 6 hpf prior to the emergence of vascular endothelial cells, as above, and analyzed expression of dsRed in GFP expressing vascular endothelium by confocal imaging of the trunk region at 72 hpf (Figure 9 A, B). The greatest number of dsRed, GFP double positive cells was observed in the posterior cardinal vein (PCV), supraintestinal artery (SIA) and subintestinal vein (SIV) (16.7, 7.3 and 19.6 cells per embryo, respectively). A subset of cells in venous Intersegmental Vessels (vISV) (Figure 9 B, arrow) were also labeled (5.8 cell per embryo). In contrast, very few labeled cells were observed in the Dorsal Aorta (DA) (0.1 cell per embryo) or arterial Intersegmental Vessels (aISV) (0.3 cell per embryo). *hand2*-derived dsRed cells were not detected in the anterior head vasculature, as expected given *hand2* expressing cells are restricted to the posterior lateral plate mesoderm in the early embryo ^12^. These results show *hand2:CreERT2* labels largely venous or vein-derived blood vessels, including the SIA, which originates partly from the PCV and partly by direct migration of late-forming endothelial progenitors ^32–35^.

**Figure 9.**
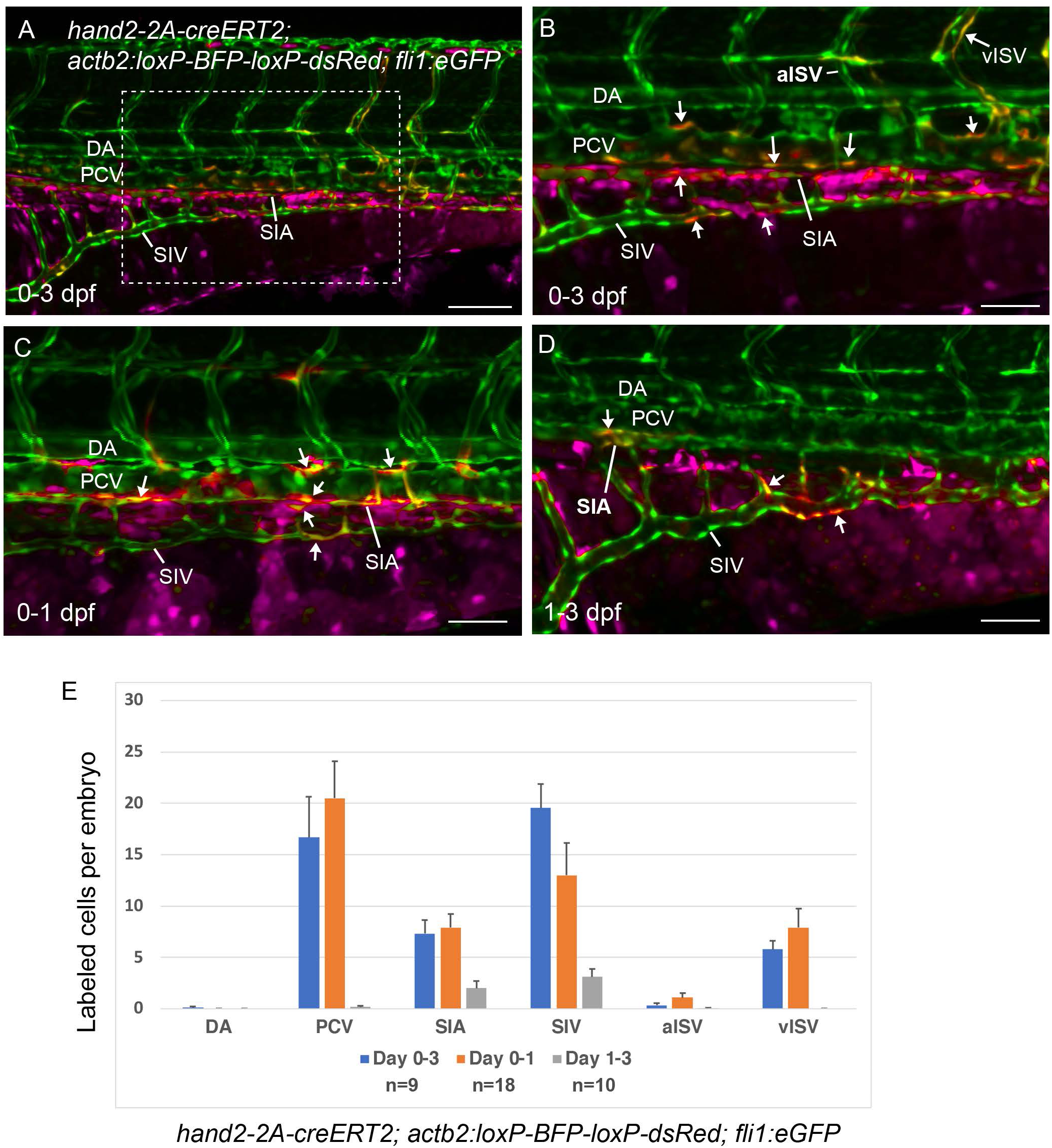
*hand2:creERT2* lineage tracing reveals contribution to the venous and intestinal vasculature. **A-D** Live embryo confocal z-stack projections of vasculature in *hand2:creERT2*; *actb2:loxP-BFP-loxP-dsRed*; *fli1:GFP* 3 dpf larvae. **A, B** Images of 3 dpf larva treated with 4- OHT beginning at 6 dpf. **A** Lateral view of larval trunk showing eGFP expression throughout the vasculature and eGFP, dsRed double positive cells (B, arrows) in the posterior cardinal vein (PCV), venous Intersegmental Vessels (vISV), supraIntestinal Artery (SIA) and subIntestinal Vein (SIV). dsRed was not detected in vascular cells in the Dorsal Aorta (DA) or aortic Intersegmental Vessels (aISV). **B** Higher magnification of boxed region shown in A. **C** Images of 3 dpf larva treated with 4-OHT from 6 hpf to 1 dpf showing eGFP, dsRed double positive cells (arrows) in the PCV, vISV, SIA and SIV. **D** Lateral view of trunk vasculature in larva treated with 4-OHT beginning at 24 hpf through 3 dpd showing eGFP, dsRed double positive cells (arrows) in the SIA and SIV. **E** Quantification of eGFP, dsRed double positive vascular endothelial cells in *hand2:creERT2*; *actb2:loxP-BFP-loxP-dsRed*; *fli1:GFP* embryos treated with 4-OHT from 6 hpf – 3 dpf (n=9), from 6 hpf – 1 dpf (n=18), and from 1 dpf – 3 dpf (n=10). Data plots show mean ± s.e.m. Scale bars: A 100 μm; B, C, D 50 μm.

Vascular endothelial progenitors emerge from the lateral plate mesoderm during early and late somitogenesis stages and coalesce into the axial vessels, the DA and PCV ^36–40^. While the initial vasculogenesis is largely complete by 24 hpf stage, when blood circulation initiates in zebrafish embryos, a recent study shows that late forming vascular progenitors continue to emerge from the secondary vascular field (SVF) along the yolk extension at 24-48 hpf, and subsequently contribute to the PCV, SIA and SIV ^35^. To determine whether both early-stage lateral plate mesoderm progenitors and late forming progenitors contain *hand2* expressing populations. We treated *hand2:CreERT2; actb2:loxP-BFP-loxP-dsRed; fli1:GFP* embryos with 4-OHT starting at 6 hpf shield stage, washing out 4-OHT at 22 hpf, followed by imaging the trunk vasculature in larva at 3 dpf (Figure 9 C). Similar to continuous 6 hpf – 3 dfp 4-OHT treatment described above, we observed dsRed, eGFP positive cells in the PCV, SIV and SIA (20.5, 13, and 7.9 cells per embryo, respectively), and in the venous ISVs (7.9 cells / embryo). Only a small number of labeled cells were located in the arterial ISVs (1.1 cells / embryo) and no labeled cells were located in the DA. These results were consistent with early lateral plate mesoderm *hand2* progenitors contributing to all venous vasculature.

To test whether there are *hand2* expressing late forming vascular progenitors that contribute to the venous vasculature after 24 hpf, we treated *hand2:CreERT2; actb2:loxP-BFP-loxP-dsRed; fli1:GFP* embryos with 4-OHT starting at 22 hpf, and analyzed the vascular contribution of switched cells at 3 dpf (Figure 9 D). The number of dsRed, eGFP positive cells was greatly reduced and limited to the SIV (3.1 cells / embryo), SIA (2 cells / embryo) and PCV (0.17 cells / embryo). Comparison of the number of dsRed, eGFP positive cells in the larval trunk vascular compartments confirmed that the early lateral plate mesoderm *hand2* population gives rise to cells in all posterior venous vessels, while late forming *hand2* vascular progenitors that arise after 24 hpf contribute to a smaller number of cells in the PCV and intestinal venous vasculature (Figure 9 E). These experiments revealed two time points at which *hand2* progenitors contribute to the venous vasculature; early progenitors present in the shield stage lateral plate mesoderm, and late emerging progenitors in the 24-48 hpf embryo. Switching at 3 dpf did not result in labeling of the vasculature, which indicates hand2 progenitor contribution to the vasculature is complete by 72 hpf. In addition to mesothelial labeling in the heart, a small number of cells were labeled in the pectoral fins and ventral fin fold, as described above and consistent with a recent publication describing *hand2* contributions to the median fin fold ^26^.

In summary, the *hand2-2A-cre* and *hand2-2A-creERT2* lines revealed the spatial and temporal contribution of *hand2* progenitors to embryonic and larval development. The results indicate a role for *hand2* transcriptional regulation in cell fates derived from the three germ layers and the neural crest, and support further investigation of the functional requirement for *hand2* in late embryo vascular development and larval organogenesis.

## Discussion

In this study we describe the isolation and characterization of Cre and CreERT2 recombinase knock-ins in the critical developmental regulator gene *hand2* in zebrafish. Using our GeneWeld CRISPR-Cas9 homology-directed repair strategy we recovered in-frame integrations located at the 3’ end of the *hand2* coding sequence. Cre and Tamoxifen regulated CreERT2 recombinase activity were validated in the expected *hand2* mesodermal and neural crest derived lineages using live embryo and larval confocal imaging with ubiquitous floxed GFP and GFP-to-mCherry switch reporter lines. Lineage tracing also confirmed *hand2* progenitors contribute to the posterior cardinal vein and venous intersegmental vessels, indicating a mesodermal origin for the venous vasculature which was predicted from previous genetic analyses, but had not been confirmed through lineage tracing or gene expression. The ability to temporally activate CreERT2 recombinase at a defined developmental stage allowed identification of a window between 24 and 72 hpf when late emerging *hand2* progenitors contribute to the intestinal vasculature. Induction of CreERT2 in the 3 dpf larva revealed *hand2* progenitors were restricted to mesodermal-derived cells in the pharyngeal arches, heart, pectoral fin, and liver, and neural crest-derived enteric neurons. The results illustrate the power of tamoxifen-regulated CreERT2 for refined temporal lineage analysis that can lead to novel discoveries of gene contributions to development. Our study emphasizes the advantage of endogenous driver knock-ins to provide comprehensive readouts of gene activity, in comparison to potentially incomplete promoter and enhancer transgenic reporters and lineage conclusions based on reporter persistence alone. The novel findings reported here support a role for *hand2* in the developing venous vasculature and during larval organogenesis in the post embryonic branchial arches, heart, liver, and developing enteric nervous system.

Precise, in-frame *hand2-2A-cre* and *hand2-2A-creERT2* integrations with linked lens-specific secondary markers were recovered at frequencies of 5% (2/43 screened; 2/10 transmitting) and 2% (1/46 screened; 1/5 transmitting), respectively, similar to our previous reports on GeneWeld precision targeted integration alleles ^1,3,4^. PCR junction and sequence analysis indicated seamless integration of the Cre cassettes into the *hand2* 3’ end target site, which was confirmed by genomic Southern blot analysis. Additional high molecular weight bands on the *hand2-2A-creERT2* Southern suggest a possible linked off target integration.

However, Cre recombinase activity in both *hand2-2A-cre* and *hand2-2A-creERT2* lines showed similar lineage tracing activity and recapitulated known patterns of *hand*2 expression in the embryo. RT-qPCR indicated expression or stability of the integration allele mRNA was reduced, but this may reflect the different primer pairs used to amplify the wildtype *hand2* vs the *hand2-2A-cre* or *-creERT2* transcripts or efficiency of reverse transcription. Regardless, neither *hand2-2A-cre* nor *hand2-2A-creERT2* allele negatively affected embryonic development and both showed homozygous viability into adulthood.

*hand2* Cre and CreERT2 recombinase activity at shield stage led to labeling of numerous expected lineages derived from the mesoderm and ectoderm. Specific lineages include the ectoderm neural crest-derived enteric neurons, as well as the mesoderm-derived posterior cardinal vein and intersegmental vessels, intestinal smooth muscle, and tissues such as heart myocardium, epicardium, bulbus arteriosus-contributing smooth muscle, pectoral fin bud and PAFF fibroblasts, and mesothelium. These observations extend the previous identification of hand2 as a marker of mesothelial progenitors in the post-gastrula lateral plate mesoderm ^18^ and enteric neurons derived from the ectodermal neural crest ^5,9^.

Notably, our work reveals that *hand2* expressing cells contribute vessels of the posterior venous vasculature, including the PCV and PCV-derived venous intersegmental vessels, the subintestinal aorta, and the subintestinal vein. Early vascular endothelial progenitors of the major trunk vasculature are known to originate in two stages. The earliest progenitors originate close to the midline and give rise to the dorsal aorta, while venous progenitors emerge more laterally and coalesce into the posterior cardinal vein ^39^. It has been previously demonstrated that *hand2* expression during these stages of vasculogenesis is localized lateral to and immediately adjacent to the venous progenitors ^12^. Furthermore, loss of *hand2* results in greatly reduced venous differentiation ^12^. Our results indicate a subset of *hand2*-expressing cells in the lateral plate mesoderm give rise to venous progenitors prior to 24 hpf stage, which helps to explain how *hand2* deficiency affects venous differentiation.

A recent study demonstrated that vasculogenesis continues past the 24 hpf stage in zebrafish embryos ^35^. The late-forming vascular progenitors originate in the secondary vascular field (SVF) along the yolk extension, and largely contribute to the PCV and intestinal vasculature, SIA and SIV during 24-48 hpf stages. Our current results show that *hand2-2A-creERT2* labeled cells contribute to the SIV, SIA and PCV at 24-72 hpf. These results strongly suggest that a subset of *hand2-*positive cells in the lateral plate mesoderm give rise to the late-forming vascular progenitors (SVF cells), observed at 24-48 hpf, and these cells then contribute to the PCV and intestinal vasculature, the SIA and SIV. These new observations illustrate the power of Cre and CreERT2 knockin lines for refined analysis of cell lineage and gene specific contributions to cell fate and development.

Taken together, the *hand2* Cre and CreERT2 drivers described in this study provide critical new tools to investigate *hand2* function and lineages in embryonic development and in post-embryonic tissue and organ morphogenesis. Our work underscores the importance of generating cell type-specific reporters that recapitulate endogenous gene expression, providing robust Cre recombinase activity with increased specificity for lineage analysis and functional gene studies.

### Experimental Procedures

#### Zebrafish care and husbandry

Zebrafish (Danio rerio) were maintained on an Aquaneering aquaculture system at 27°C on a 14 h light/10 h dark cycle. The wildtype WIK strain used to generate targeted integration lines was obtained from the Zebrafish International Resource Center (https://zebra fish.org/home/guide.php). Embryos were collected and maintained at 28.5°C in E3 embryo media ^41^ and staged according to published guidelines ^42^. Transgenic reporter lines used in this study were *Tg(-3.5ubb:loxP-EGFP-loxP-mCherry)cz1701* ^28^ (ubi:Switch), *Tg(–1.5hsp70l:loxP-STOP-loxP-EGFP,cryaa:Venus)zh701* ^29^ (hsp70l:Switch), *Tg(3.5ubi:loxP-lacZ-loxP-eGFP)cn2* ^25^ (ubi:Hulk), *Tg(fli1:GFP)*y1 ^43^ (*fli1:GFP*), and *Tg(actb2:loxP-BFP-loxP-dsRed)sd27* ^44^.

Experimental protocols used in this study were approved by the Iowa State University Institutional Animal Care and Use Committee (IACUC-20-058, IBC-20-071) in compliance with American Veterinary Medical Association and the National Institutes of Health guidelines for the humane use of laboratory animals in research.

#### Isolation of *hand2-2A-cre* and *hand2-2A-creERT2* lines using GeneWeld CRISPR-Cas9 targeted integration

*2A-cre,gcry1:mRFP* and *2A-creERT2; gcry1:eGFP* cassettes were integrated in frame at a CRISPR-Cas9 gRNA site located at the 3’ end of the *hand2* coding sequence in exon 2 using GeneWeld targeted integration as previously described ^1–3^. Cas9 capped mRNA was *in vitro* synthesized from expression vector pT3TS-nls-Cas9-nls^45^ (Addgene #46757) linearized with XbaI (New England Biolabs, R0145S) using the Ambion mMessage Machine T3 Transcription Kit (Thermo Fisher, AM1348). *in vitro* synthesized mRNA was purified with the RNA Clean and Concentrator Kit RCC (Zymo, R1013). The *hand2* gRNA 5’-GCCACAGCATGTTTGGGCAT-3’ was tested by co-injection of 25 pg gRNA and 300 pg Cas9 mRNA into WIK wildtype embryos. Genomic DNA used for PCR was extracted from 1 dpf injected embryos by incubation in 50mM NaOH for 30’ at 95°C and neutralized by addition of 1/10th volume 1 M Tris pH 8.0. Amplicons were Sanger sequenced and mutagenesis efficiency was evaluated using Synthego ICE software https://www.synthego.com/products/bioinformatics/crispr-analysis (Synthego). 5’ and 3’ homology arms (Table 2) were assembled in the GeneWeld vectors *pPRISM-2A-cre,gcry1:mRFP* (Addgene 117790) and *pPRISM-2A-cre,gcry1:eGFP* (Addgene 117791) as described ^2^. The 5’ homology arm also contained the remaining 17 nucleotides of the *hand2* coding sequence located downstream of the CRISPR-Cas9 cut site, with scrambled codons to prevent cutting by the *hand2* genomic gRNA. The *cre* cDNA in the *hand2*-homology arm assembled-*pPRISM-2A-cre,gcry1:eGFP* targeting vector was replaced with *creERT2* from pENTR_D_creERT2^28^ (Addgene 27321) using *creERT2* PCR and vector restriction enzyme digestion fragments in a New England Biolab Hi-Fi cloning (NEB #E2621S) reaction.

**Table 2.**
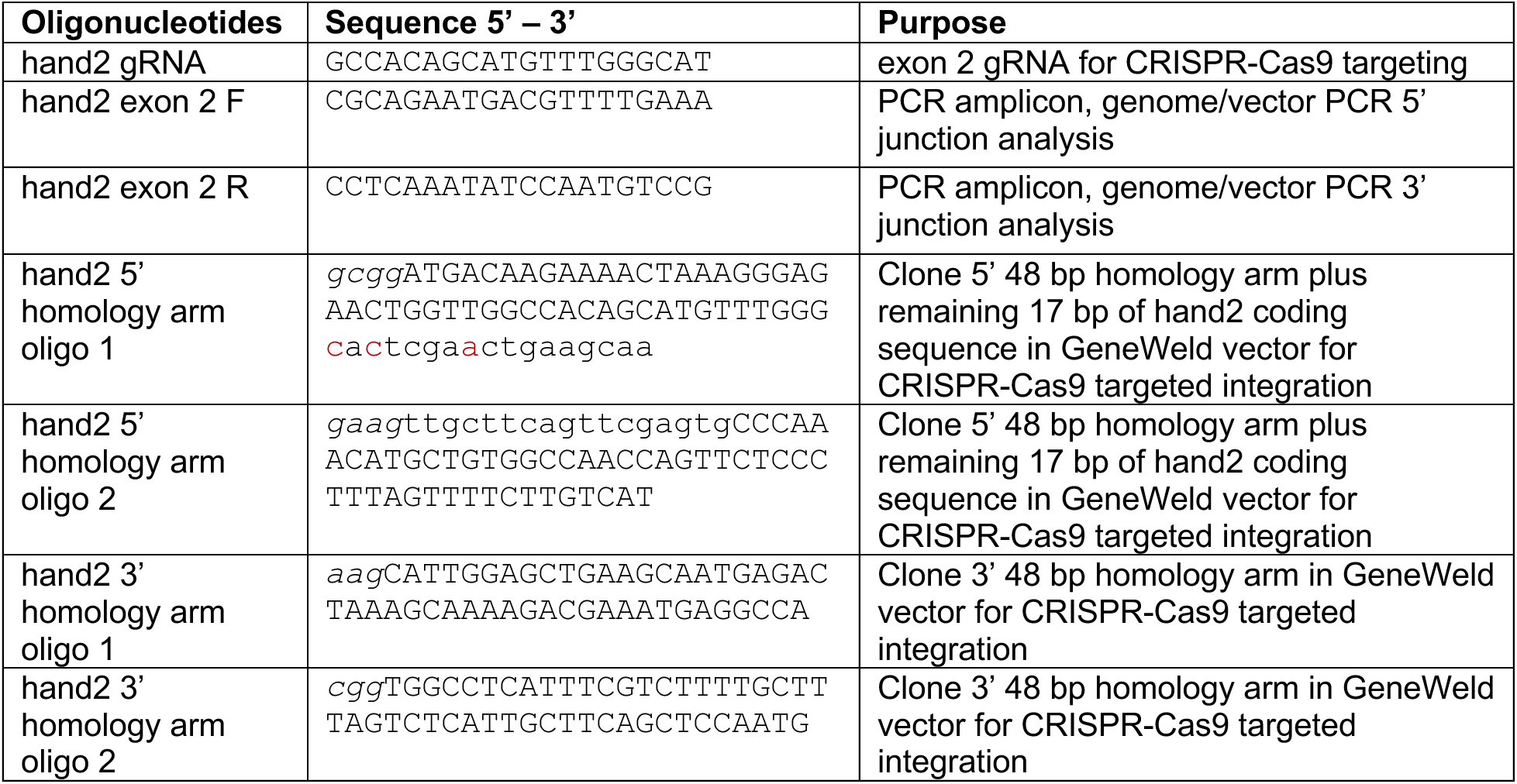
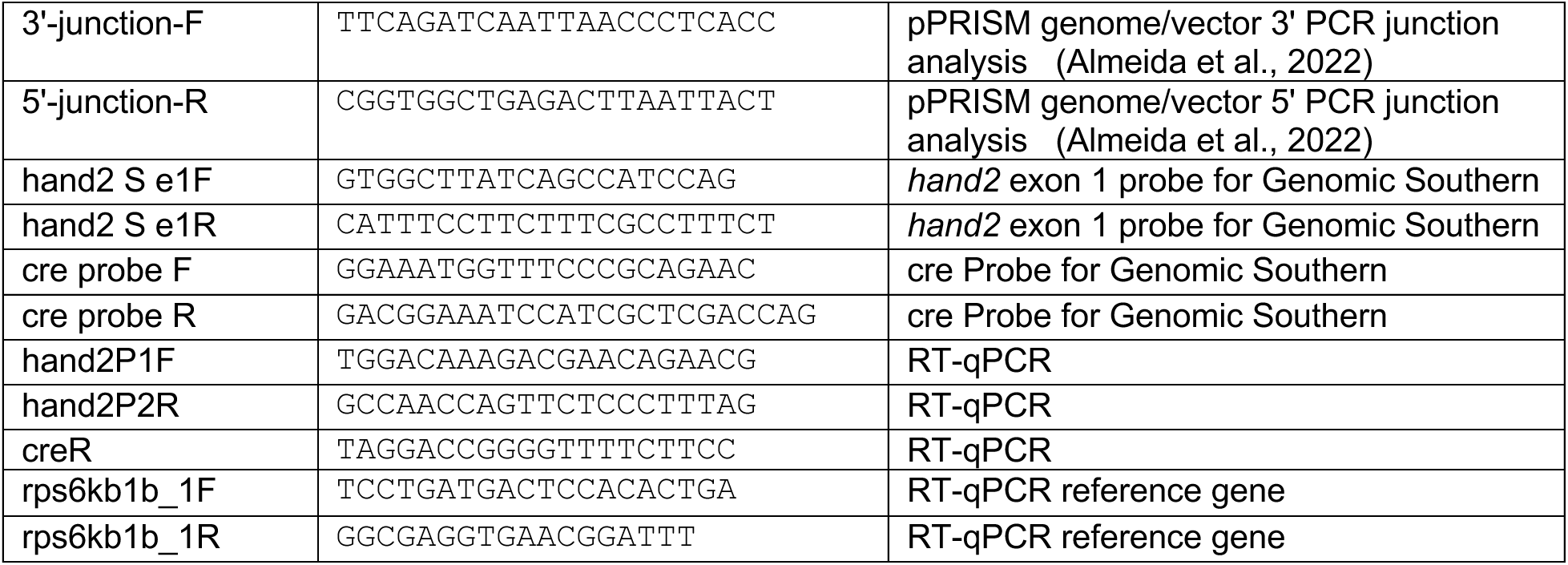
Sequences of oligonucleotides used in this study. Oligonucleotide Sequences of *hand2* 3’ target site gRNA, primers used for PCR and RT-qPCR, and oligonucleotides to build *hand2* 48 bp homology arm pPRISM-2A-cre and-2A-creERT2 targeting vectors. Sequences in lowercase replace the end of the *hand2* coding sequence with scrambled codons (shown in red) to prevent the genomic gRNA from cutting in the pPRISM-cre vector 5’ homology arm. Sequences in lowercase italics are sequences complementary to BfuAI and BspQI overhang after restriction enzyme digestion of the GeneWeld pPRISM-2A-cre and-2A-creERT2 vectors.

To isolate *hand2-2A-cre, gcry1:mRFP* and *hand2-2A-creERT2, gcry1:eGFP* knock-in alleles, 2nl volume containing 25pg *hand2* gRNA, 25pg Universal gRNA, 10 pg targeting vector, and 300pg Cas9 mRNA were coinjected into wildtype WIK 1 cell stage embryos. 3 dpf larvae expressing the linked lens specific *gcry1:mRFP* or *gcry1:eGFP* secondary markers were selected for isolation of genomic DNA and PCR based 5’ and 3’ genomic DNA-integrated cassette junction analysis. 50 injected sibling embryos positive for lens specific fluorescence secondary markers were raised to adulthood. Potential founder adults were screened by crossing to the ubi:Switch line and examining embryos for the expected secondary marker and for GFP to mCherry switching in the heart. F1 embryos were tested for precise 5’ and 3’ junctions by PCR and Sanger sequencing. Sibling embryos from positive founders were raised to adulthood and fin clipped to confirm the presence of the precise integration allele. Positive F1 adults were outcrossed to wild type WIK, and single positive F2 adults were used to establish independent transgenic lines.

#### RT-qPCR

Experiments to measure relative transcript levels by Reverse Transcription-quantitative PCR were designed and performed according to MIQE and updated guidelines (<u>Bustin et al.,</u> <u>2009</u>; <u>Taylor et al., 2019</u>). Three biological replicates were performed, with each replicate representing embryos from a different mating pair of heterozygous F2 adults. 40 randomly selected 4 dpf larvae from incrosses of heterozygous *hand2-2A-cre,gcry1:mRFP/+* or *hand2-2A-creERT2; gcry1:eGFP/+* were collected for RNA extraction and genotyping. Individual dissected head tissue was placed in DNA/RNA shield (Zymo Research R1100-50) and individual trunk tissue was placed in 50 mM NaOH for genotyping. Five heads of each genotype were pooled and total RNA extracted using the Direct-zol RNA Microprep kit (Zymo Research, R2060) and quality determined using a ThermoScientific Nanodrop 2000 spectrophotometer. RNA samples with a A260/A280 >1.90 were normalized to the same concentration. Standard curve was constructed by performing one-step RT-qPCR (Luna® Universal One-Step RT-qPCR Kit, NEB R3005) using either heterozygous *hand2-2A-cre,gcry1:mRFP/+* or wild type larvae.

Standard curve reaction of 40 cycles of amplification was run at 60°C annealing temperature using 400 nM primer concentration and five starting RNA amounts per reaction (100ng, 20ng, 4 ng, 0.8ng, and 0.16ng) with two replicates per condition. A single sharp melt curve peaked was obtained for each primer set at the desired mRNA quantity range and subsequence agarose gel electrophoresis revealed a single amplicon of intended size with each primer set. Primer efficiency was calculated as described (Bustin et al., 2009; Taylor et al., 2019) and primer pairs with 86–102% efficiency were used for RT-qPCR control and test samples.

RT-qPCR was performed on each sample in triplicate using Luna® Universal One-Step RT-qPCR Kit (NEB R3005) on a CFX Connect Real-Time System (Bio-Rad). 100ng of total RNA per reaction was used for *hand2-2A-cre,gcry1:mRFP* RT-qPCR experiments. 60ng of total RNA per reaction was used for *hand2-2A-creERT2; gcry1:eGFP* RT-qPCR experiments.

Relative gene expression was calculated using the Pfaffl Method which considers amplification efficiency differences among different primer pairs. *hand2*, *hand2-2A-cre* and *hand2-2AcreERT2* levels were measured relative to reference gene *rps6kb1b*. The *hand2* forward primer was located in exon 1, the *hand2* reverse primer was located in exon 2 coding sequence upstream of the translation stop codon, and the *cre* reverse primer was located in the *cre* coding sequences shared between *cre* and *creERT2* cDNAs. RT-qPCR primer sequences are listed in Table 2.

#### WGS and analysis

Whole Genome Sequencing was performed with PacBio Long Red Sequencing, 30X coverage, followed by de novo contig assembly and analysis (MedGenome, Inc). PacBio libraries were prepared with genomic DNA extracted from 60mg muscle tissue from *hand2-2A- cre, gcry1:mRFP^is^*^78*/+*^ and *hand2-2A-creERT2, gcry1:eGFP^is^*^79*/+*^ adults and flash frozen in liquid nitrogen. *hand2-2A-cre* and *hand2-2A-creERT2* donor plasmid sequences were mapped to the assembled contigs using GeneMap, contigs were mapped to the Zebrafish genome assembly GRCz11 at the UCSC genome browser.

#### 4-OHT treatment and imaging

*hand2-2A-creERT2*; *ubi:switch* embryos were treated continuously with 10 μM 4- hydroxytamoxifen (4-OHT) (Sigma H6278) in E3 embryo media from 6 hpf until imaging at 2, 3 and 5 dpf. *hand2-2A-creERT2*; *ubi:switch* larvae were treated with 5 μM or 6.7 uM 4- hydroxytamoxifen beginning at 3 dpf and imaged at 5 dpf. *hand2-2A-creERT2*; *hsp70l:loxP-STOP-loxP-EGFP* embryos were heat shocked for 1 hour at 37°C and imaged 3 hours later as described ^30^. *hand2-2A-creERT2*; *actb2:loxP-BFP-loxP-dsRed* were treated with 10 μM 4-OHT from 6 hpf through 3 dpf, from 6 hpf through 1 dpf, or from 1 dpf through 3 dpf, followed by live imaging at 3dpf. For live imaging embryos were incubated in E3 embryo media ^41^ containing 0.003% 1-phenyl 2-thiourea (PTU) (Sigma P7629) to inhibit pigment synthesis. Embryos and larvae were anesthetized in 0.015% Tricaine MS-222 Ethyl 3-aminobenzoate methanesulfonate (Sigma E10521) and mounted in 1.2% low melt agarose (Promega V2111) in embryo media containing 0.003% PTU/0.015% Tricaine. For vibratome sectioning, zebrafish embryos were harvested at the stages indicated and fixed in 4% PFA at 4°C overnight (n= 8 embryos/stage). Yolk was manually removed from the embryos using forceps before embedding in 4% low-melting agarose. Agarose blocks were chilled at 4°C for a minimum of 2 hours. 150μm transverse sections were cut on a vibratome (Leica). Sections were transferred to 24-well tissue culture plate and stained with DAPI (1μm/ml) and Alexa 647-Phalloidin (1:100) in PBDT (PBS, 0.1% Triton X-100, 1% DMSO) for 1 hour.

Live embryo and larvae fluorescence and bright field images were captured using a Zeiss AxioZoomV16 with Apotome, a Zeiss LSM 800 laser scanning confocal microscope, a Zeiss LSM 880 laser scanning confocal microscope, and a Nikon Eclipse confocal microscope. Nikon Elements Denoise.ai algorithm was used to remove noise from Nikon Eclipse confocal images of larval vasculature with high background signal. Maximum intensity projections of Nikon Eclipse confocal Z-stacks of larval vasculature were obtained using Fiji (Image J) software. Double positive *fli1:GFP* and dsRed cells in the vasculature Z-stack images were counted manually. 5 dpf larva were fixed in 3.7% formaldehyde and imaged on a Nikon Ti-E Inverted Microscope, A1R Confocal, GaAsp PMTs, Andor iXon 888 EMCCD Widefield Camera (Nikon Instruments Inc, Melville, NY). Post image processing and data analyses were conducted using Imaris (Bitplane, Concord, MA).

## Quantification and Statistical Analysis

Excel software was used to generate data plots showing mean ± s.e.m.

## Data Availability

DNA constructs and transgenic zebrafish lines generated in this study are available on request to M. McGrail.

## Acknowledgements

The authors thank Dr. Pascal Lafontant, Grinnell College, for discussions and Dr. Juan Manuel Gonzalez Rosa, Boston College, for the *ubi:Hulk* reporter line. Graphical figures were Created with BioRender.com with a paid Biorender license. This work was supported by NIH ORIP R24 OD020166 (MM, JJE, IF, KJC, SCE); NIH NIGMS 1T32GM141742-01 and NIH NHLBI F31 HL167580 (HRM); NIH NHLBI K99 1K99HL168148–01 (RLL); the Anschutz Medical Campus, the Children’s Hospital Colorado Foundation, and NIH NHLBI R01 HL168097-01A1 (CM); CM holds The Helen and Arthur E. Johnson Chair for the Cardiac Research Director; NIH NIDDK R01 DK117266-01A1 (CY), the Peter and Tommy Colucci Award for PFIC Research (CY), and NIH NHLBI R01 HL153005 (SS).

## Ethics approval statement

Experimental protocols used in this study were approved by the Iowa State University Institutional Animal Care and Use Committee (IACUC-20-058, IBC-20-071) and the CU Anschutz Institutional Animal Care and Use Committee (979), in compliance with American Veterinary Medical Association and the National Institutes of Health guidelines for the humane use of laboratory animals in research.

